# ClickArr: a novel, high throughput assay for evaluating β-arrestin isoform recruitment

**DOI:** 10.1101/2022.09.28.509965

**Authors:** Alexander R. French, Yazan J. Meqbil, Richard M. van Rijn

## Abstract

Modern methods for quantifying signaling bias at GPCRs rely on using a single β-arrestin isoform. However, it is increasingly appreciated that the two β-arrestin isoforms have unique roles, requiring the ability to assess β-arrestin isoform preference. Herein, we present ClickArr, a live-cell assay that simultaneously reports recruitment of both β-arrestin isoforms as they compete for interaction with GPCRs. We demonstrate that an agonist can have β-arrestin isoform bias, potentially opening up a new dimension for drug development.

---

Biased signaling at GPCRs has been a focal point of drug design in recent years, with the discovery that “biased” agonists can direct cell signaling through one pathway over another, promising to produce next-generation pharmaceuticals that have established or new therapeutic effects while lacking deleterious “side effects” from the activation of undesirable pathways^1–4^. Most biased signaling efforts to date have focused on the relative signaling of β-arrestins versus G proteins^1,2^. However, it is commonly found that β-arrestin isoforms can play distinct and often opposing roles in disease models^5^ including rheumatoid arthritis^6^ and Parkinson’s disease^7^, as well as in modulating the effects of addictive substances such as amphetamine^8^ and psychedelic substances such as lysergic acid diethylamide (LSD)^9^. At the molecular level the two isoforms play different roles in regulating the delta opioid receptor (δOR), a GPCR involved in pain and mood regulation^10^. Similarly, the two isoforms have differential effects on regulating δOR-induced anxiolytic and fear-reducing effects^11^.

To take full advantage of the therapeutic potential of GPCRs, biased agonists would ideally account for differences in β-arrestin isoform signaling. In cells, the two β-arrestin isoforms compete for a single site at a GPCR. In currently available drug screening platforms, a single isoform is overexpressed and this competition not accounted for. Moreover, comparing activity at each isoform requires normalizing across different tests and even cell lines. Thus, comparing the performance of both isoforms doubles the time and material costs of the study, discouraging it in common practice.

Here we present ClickArr, a <u>click</u> beetle luciferase-based assay for <u>arr</u>estin recruitment that allows simultaneous readout of both β-arrestin 1 and 2 from the same cell. ClickArr is optimized to run in multiwell plate format for fast evaluation of lead compounds and arrestin signaling characteristics. Our design for ClickArr takes advantage of the bright click beetle green (CBG) and click beetle red (CBR) luciferases, which both catalyze the oxidation of the same substrate, D-luciferin, but emit light at different wavelengths^12,13^. It was previously demonstrated that the N-terminal fragment of both CBG and CBR can complement with a C-terminal fragment from CBG and retain the spectral properties of the full-length proteins^14^. We reasoned that fusing a C-terminal fragment of CBG to a GPCR, and then fusing N-terminal fragments of CBG and CBR to β-arrestin 1 and β-arrestin 2, respectively would allow us to monitor the recruitment of each isoform as they compete for GPCR binding sites in a cell (Fig. 1a).

**Fig. 1.**
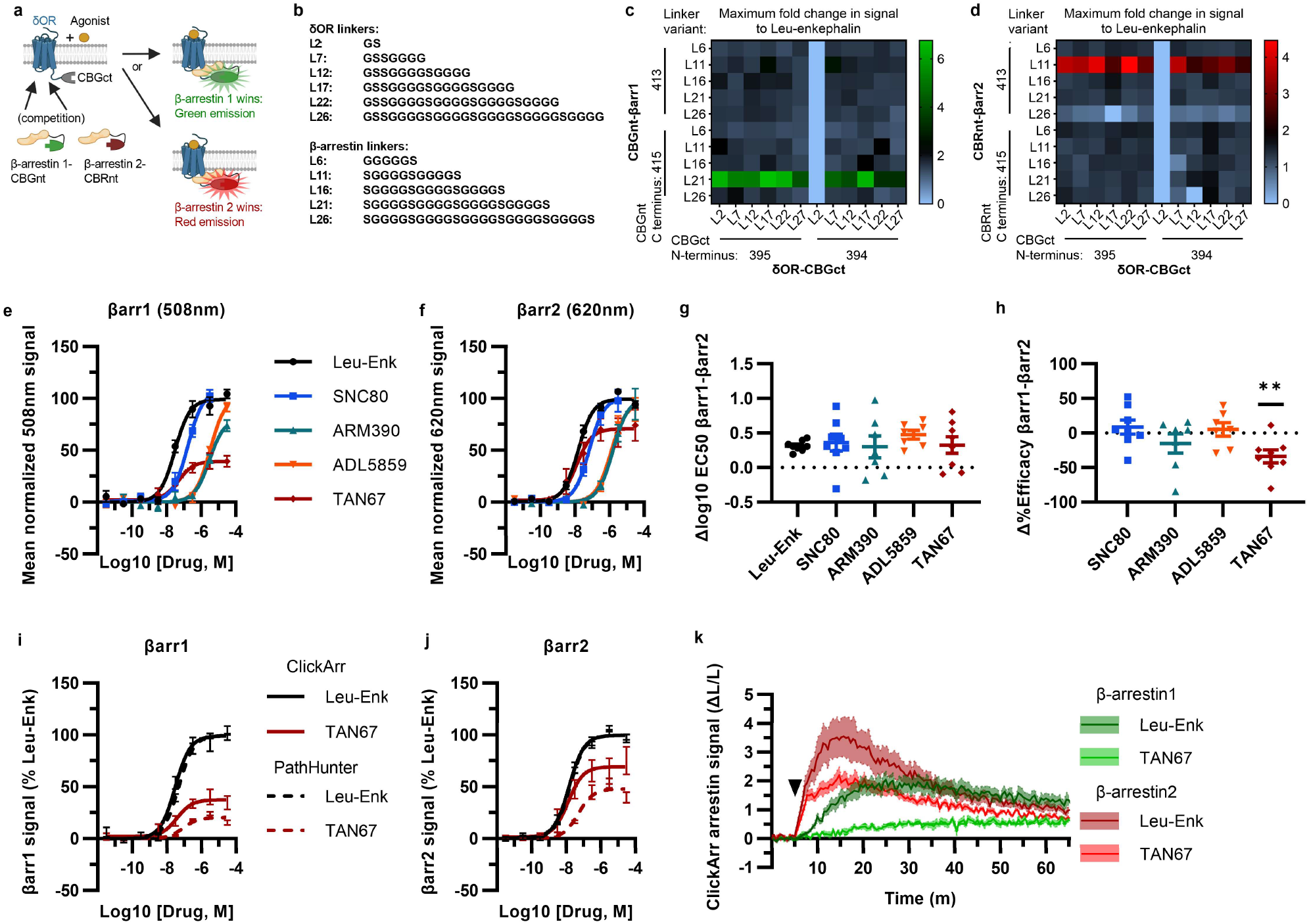
Development of ClickArr assay using the δOR. **a**, Schematic showing assay design enabling simultaneous, independent readout of β-arrestin recruitment to the δOR. CBGnt and CBGct: N- and C-terminal fragment of click beetle green luciferase, respectively. CBRnt: N-terminal fragment of click beetle red luciferase. **b**, Linker sequences screened between luciferase fragment and the δOR (top) or β-arrestins (βarr) (bottom). **c**,**d**, Maximum fold response to the endogenous δOR ligand leu-enkephalin for β-arrestin 1 (βarr1) (**c**) and β-arrestin 2 (βarr2) (**d**) constructs with various linkers and luciferase fragment termini. **e**,**f**, ClickArr assay results using top screen hits from **c** and **d** showing arrestin recruitment for the βarr1 (**e**) and βarr2 (**f**). **g**, Agonists tested do not show a significant shift in potency relative to reference agonist leu-enkephalin. **h**, However, TAN67 shows significantly lower efficacy for βarr1 than βarr2. **, p < 0.01 relative to Δ%Efficacy for leu-enkephalin (defined as zero). **i**,**j**, ClickArr curves for leu-enkephalin (Leu-Enk) and TAN67 compare well to the PathHunter assay, the state-of-the-art assay for β-arrestin recruitment. **k**, ClickArr assay enables kinetic analysis of β-arrestin isoforms by different agonists, where bias measured at equilibrium is reflected in the time courses of recruitment (Figure S2). Curves are mean; shaded area ± SEM, n = 6.

Given the established roles of β-arrestin 1 and 2 at the δOR^10,15,16^, and the availability of agonists ranging from partial recruiters to super-recruiters relative to the endogenous agonist leu-enkephalin^15,16^, we pursued this receptor in our initial proof-of-concept as a meaningful testcase for finding agonists biased toward a β-arrestin isoform. We adopted a linker screen previously used for optimizing split click beetle luciferase complementation assays^17^ and generated a small library of δOR and β-arrestin constructs fused to fragments of click beetle luciferase (Fig. 1b). We evaluated two termini for each click beetle luciferase fragment used. We optimized the ClickArr assay to be carried out in 384-well format, allowing us to perform a full doseresponse curve of each pair of constructs to leu-enkephalin. Plotting the maximum fold change in signal to leu-enkephalin revealed that most combinations produced at least a 50% increase in signal for β-arrestin 1 and 30% for β-arrestin 2 suggesting a robust assay design (Fig. 1c,d). Interestingly, for each β-arrestin isoform there was a critical linker/fragment terminus combination that provided a much higher response than the others. In contrast, the signal change was less dependent on the δOR fusion construct used, with the exception of one construct. Given these results, δOR-L22-CBGct^395^, βarr1-L21-CBGnt^415^, and βarr2-L11-CBRnt^413^ were carried forward as the “ClickArr” assay (Fig. 1c,d).

Class A GPCRs such as the δOR recruit β-arrestin 2 more strongly than β-arrestin 1, demonstrating that some receptors can distinguish between β-arrestin isoforms^18^. This raised to us the possibility that different receptor:agonist complexes could have a relative bias in β-arrestin isoform recruitment. To establish whether ClickArr could detect differences in β-arrestin isoform recruitment, we characterized our reference compound leu-enkephalin in parallel with a small panel of tool agonists (Fig. 1e,f; Table S1). The rank order in potency for the agonists was the same for β-arrestin 1 and 2 recruitment were similar (Fig. 1g). However, we found a significant shift in the efficacy values for TAN67 relative to the endogenous agonist leu-enkephalin (one sample t test t = 3.663, p = 0.008), with TAN67 recruiting β-arrestin 2 with higher efficacy than β-arrestin 1 (Fig. 1h). This result establishes the ability of GPCRs to bias recruitment between β-arrestin isoforms depending on the agonist bound.

The PathHunter assay is a commercial assay for high-throughput evaluation of recruitment of a single β-arrestin isoform to a receptor and is a standard bearer in this field^19^. To compare the performance of ClickArr to the PathHunter assay, and independently confirm the presence of β-arrestin isoform bias at the δOR, we ran the δOR: β-arrestin 1 and δOR: β-arrestin 2 PathHunter assays. We found that the performance of leu-enkephalin is nearly identical in the ClickArr and PathHunter assays (Fig. 1i,j; Supplementary Table 1). Similarly, though the ClickArr assay trends toward being more sensitive to the partial agonist TAN67, this difference was not significant (Supplementary Fig. 2). Importantly, the PathHunter assay confirmed the presence of an efficacy bias of TAN67 for β-arrestin 2 recruitment over β-arrestin 1 (Fig. 1i,j; Supplementary Fig. 2b). Our results with the PathHunter assay therefore validate the β-arrestin isoform bias detected in the ClickArr assay and support the idea that GPCR:agonist complexes can distinguish between β-arrestin isoforms.

A unique feature of ClickArr is that it is a live-cell assay, which should enable kinetic studies of β-arrestin recruitment. To test our hypothesis, we recorded β-arrestin 1 and 2 recruitment to δORs in the presence of leu-enkephalin or TAN67 for over 1 hour. Comparing the Leu-enkephalin and TAN67 traces for each isoform, the peak β-arrestin recruitment signal of TAN67 is approximately 59% of leu-enkephalin for β-arrestin 2, but only 34% of leu-enkephalin for β-arrestin 1, suggesting isoform bias at saturating (~32 mM) agonist concentration (Fig. 1k). The multiwell plate format of ClickArr allowed us to evaluate kinetic responses to both Leu-Enk and TAN67 at multiple concentrations simultaneously (Supplementary Fig. 2). This enabled us to evaluate bias through analysis of the initial reaction velocities, which can be a more sensitive measure of bias than equilibrium or endpoint analyses^20^. The kinetic data (Supplementary Fig. 2) should in principle replicate the β-arrestin isoform bias seen in our single timepoint study (Fig. 1h), which is indeed the case.

Therefore, we have validated ClickArr as a novel assay simultaneously reporting recruitment of both β-arrestin isoforms to the δOR. We found that different agonists complexed with δOR could have different relative efficacies for recruiting the two β-arrestin isoforms (Fig. 1h). Given that β-arrestin 1 and 2 can have opposing roles at GPCRs, this finding has important implications for future drug design. Namely, ClickArr could help identify drugs with β-arrestin isoform bias that have reduced off-target pathway activation and subsequently improved sideeffect profiles. For example, at the δOR, β-arrestin 2 recruitment is associated with necessary receptor internalization and resensitization, whereas β-arrestin 1 recruitment targets δORs for degradation leading to rapid tachyphylaxis^10^. As δORs are a target for chronic indications, such as migraine and neuropathic pain, avoiding β-arrestin 1-induced tachyphylaxis would improve the repeated performance of therapeutics at this receptor^21^. In fact, TAN-67 has preclinical efficacy in reducing alcohol intake in mice^15^, and can be considered a potential treatment for alcohol use disorder, an indication that would also require chronic use. Thus, the finding that arrestin recruitment at TAN-67 is biased away from β-arrestin 1 could make this a more exciting lead candidate.

Conceivably, ClickArr could be extended to GPCRs beyond the δOR, making it possible that ClickArr could be expanded to a diverse array of prospective drug targets. Beyond providing readouts for the development of new agonists at these receptors, ClickArr may help retroactively explain instances where a study examining β-arrestin 2 recruitment alone to establish G protein vs arrestin bias failed to find a correlation between G protein bias and side effect profile. For example, if agonists that poorly recruit β-arrestin 2 still significantly recruit β-arrestin 1, an apparent G-biased agonist could still activate β-arrestin-linked pathways.

The β-arrestin isoform bias we observed for TAN67 is partial, meaning that recruitment of neither isoform is completely eliminated. Thus, the full extent to which different receptors can be biased toward a given isoform is not yet known and is likely to vary between receptors. It is not yet known whether isoform bias can be strengthened to a clinically meaningful degree. Nonetheless ClickArr has opened a new dimension of agonist bias to explore and is itself an effective, high-throughput tool for investigating these questions.

## Acknowledgements

The research reported in this publication was supported by the NIAAA and the NIMH of the NIH under Award Numbers R01AA025368 (RMvR) and F32MH115432 (ARF). We also thank the Purdue Pharmacy Imaging Core for the use of their plate reader.

## Author contributions

RMvR conceived of assay. RMvR, YJM and ARF designed constructs. ARF cloned constructs and developed assay. ARF and YJM collected data. ARF prepared manuscript. RMvR, ARF, and YJM revised manuscript.

## Methods

### Molecular cloning

#### Sequences of fragments inserted into gene vectors to construct original ClickArr fragments

/*NotI*/-δOR(res. 338-372)-Gly-Ser-CBGct(res. 395-542)-STOP-/*PasI*/

/*NheI*/-CBGnt(res. 1-413)-Ser-Gly-Leu-Lys-Ser-Arg-Arg-Ala-Leu-Asp-Ser-Ala-β-arrestin 1(res. 2-169)-/*BspEI*/

/*NheI*/-CBRnt (res. 1-413)-Ser-Gly-Leu-Lys-Ser-Arg-Arg-Ala-Leu-Asp-Ser-Ala-β-arrestin 2(res. 2-197)-/*AgeI*/

For each, the portion of either the δOR, β-arrestin 1, or β-arrestin 2 encoded was set to replace the original sequence cut out by the restriction site without altering the protein’s sequence in the final construct. The exception is in the arrestin constructs, the start codon methionine is removed, as it was no longer necessary once the N-terminal luciferase fragments were added.

#### Cloning approach

We created the original δOR-CBGct construct by digesting a pCDNA3.1 vector containing an N-terminally FLAG-tagged δOR sequence^1^, generously provided to us by the Whistler lab, with *NotI* and *PasI*. Into this we ligated a *NotI*/*PasI*-digested fragment of a custom-ordered vector containing the C-terminal (res. 395-542) click beetle green (CBG) fragment (CBGct)^2^. To generate CBGnt-β-arrestin 1, we digested a pCDNA3.1 vector encoding β-arrestin 1 (cDNA.org) with *NheI/BspEI*. Into this we ligated a NheI/BspEI-digested fragment of a custom-ordered vector encoding the N-terminal fragment (res. 1-413) of CBG (CBGnt). Similarly, to generate CBRnt-β-arrestin 2, we digested a pCDNA3.1 vector encoding β-arrestin 2 (cDNA.org) with *NheI* and *AgeI*. Into this we ligated an NheI/AgeI-digested fragment from a custom-ordered vector encoding N-terminal fragment (res. 1-413) of click beetle red (CBRnt). All ligations were performed with T4 DNA ligase (New England Biolabs) according to the manufacturer’s guidelines. Sequence maps of fragments may be found in the supplementary material.

To optimize construct performance, we screened a library of constructs having the linker sequences used by Misawa et al 2010^3^. To mutate the original constructs and generate the linker library, we used the NEBuilder HiFi DNA Assembly kit (New England Biolabs). We designed primers encoding the desired linkers and flanking regions that overlap with the sequences of the DOR, arrestins, and luciferase fragments according to the manufacturer’s guidelines. Two sets of primers were ordered for each construct to test different termini of the luciferase fragments. For DOR-CBGct, primers were ordered to include the fragments CBG(395-542) and CBG(394-542). For CBGnt-Barr1, CBG(1-413) and CBG(1-415) were tested. For CBRnt-Barr2, CBR(1-413) and CBR(1-415) were tested. The linker sequences tested were:

#### DOR-linker-CBGct

(original) GS

GSSGGGG

GSSGGGGSGGGG

GSSGGGGSGGGGSGGGG

GSSGGGGSGGGGSGGGGSGGGG

GSSGGGGSGGGGSGGGGSGGGGSGGGG

#### CBGnt-linker-Barr1 and CBRnt-linker-Barr2

(original) SGLKSRRALDSA

GGGGGS

SGGGGSGGGGS

SGGGGSGGGGSGGGGS

SGGGGSGGGGSGGGGSGGGGS

SGGGGSGGGGSGGGGSGGGGSGGGGS

### ClickArr assay

#### General protocol

A detailed protocol printout is included in the supplemental methods. Briefly, cells were transfected using XtremeGene9 reagent (Roche) according to the manufacturer’s instructions using an equal molar ratio of each construct. For the screen, carrier DNA was included in the transfections to compensate for the lack of the second β-arrestin construct, so that all the transfections had the same total DNA. Two days later, transfected cells were rinsed with DPBS (Gibco), dissociated with trypsin-EDTA (Gibco) and 15,000 – 22,000 cells seeded into wells in a 384 well plate in Opti-MEM (Gibco). Plates were sealed with Aeraseal (Millipore-Sigma) and put in a 37C incubator maintained at 5% CO2 for 30 minutes to equilibrate. A 2 mM solution of D-luciferin (GoldBio) was made in assay buffer (AB, HBSS (Gibco) + 20 mM HEPES). After equilibration, 7.5 μl of this was added to each well, the plate spun, resealed, and returned to the incubator. Agonist solutions were prepared at 4x concentration in AB and 5 μl added to the wells after 30 minutes. Plates were again spun, resealed, and returned to the incubator. A Biotek Synergy4 plate reader equipped with 508/20 and 620/10 EM filters was preheated to 37C and readings performed 30 minutes after drug addition (0.5s integration time).

#### Kinetic experiments with ClickArr

In our hands 20 wells could be read in both colors with a timestep of 30 seconds. We thus ran each kinetic assay with the indicated concentrations of agonists (four concentrations of leu-enkephalin and TAN67) in duplicate along with two vehicle wells to control for signal drift.

### Data analysis

The Synergy4 plate reader was controlled using Gen5 v2.04 software. The data was analyzed and plotted using GraphPad Prism 9 software. Dose-response curves were analyzed using the three parameter “log(agonist) vs. response” algorithm in Prism 9. The fold change data in the screen is reported as top/bottom of a single dose response curve run in duplicate for each β-arrestin:δOR pair. In order to generate a more authentic run error, the data in tables S1 and S2 represent the means of error of the fit values from the independent assays, rather than the fit error on the averaged curves in the main text.

In kinetic experiments, the signal for each well was normalized to its mean baseline over the 2 minutes preceding the drug addition and then the mean normalized signal from the vehicle wells subtracted to control for drift. We find a 5 minute baseline is sufficient to equilibrate to the plate reader.

## Supplemental information

**Supplemental Table 1:** Fit values for δOR ClickArr assay

| | β-arrestin 1 | | | β-arrestin 2 | | |
| --- | --- | --- | --- | --- | --- | --- |
|  | pEC<sub>50</sub> | EC<sub>50</sub> (nM) | E<sub>max</sub> (%) | pEC<sub>50</sub> | EC<sub>50</sub> (nM) | E<sub>max</sub> (%) |
| Leu-enkephalin | -7.49 (0.11) | 32.2 | 100 | -7.81 (0.11) | 15.7 | 100 |
| SNC80 | -6.81 (0.17) | 153.5 | 110 (7.3) | -7.15 (0.08) | 71.4 | 102 (11) |
| ARM390 | -5.53 (0.11) | 2978 | 82.4 (6.5) | -5.83 (0.08) | 1487 | 97.6 (19) |
| ADL5859 | -5.47 (0.05) | 3401 | 102 (4.7) | -5.95 (0.08) | 1134 | 97.1 (13) |
| TAN67 | -7.54 (0.29) | 28.7 | 37.2 (5.2) | -7.90 (0.19) | 12.6 | 71.6 (12) |

All values mean (SEM); pEC<sub>50</sub> values reported in log<sub>10</sub>(M). n = 7-8.

**Supplemental Table 2:** Fit values for PathHunter assay

| | β-arrestin 1 | | | β-arrestin 2 | | |
| --- | --- | --- | --- | --- | --- | --- |
|  | pEC<sub>50</sub> | EC<sub>50</sub> (nM) | E<sub>max</sub> (%) | pEC<sub>50</sub> | EC<sub>50</sub> (nM) | E<sub>max</sub> (%) |
| Leu-enkephalin | -7.30 (0.07) | 50.3 | 100 | -7.77 (0.05) | 17.0 | 100 |
| TAN67 | -7.08 (0.07) | 83.9 | 20.4 (1.4) | -7.30 (0.04) | 51.9 | 48.4 (5.2) |

All values mean (SEM); pEC<sub>50</sub> values reported in log<sub>10</sub>(M). n = 5.

**Supplemental Fig. 1.**
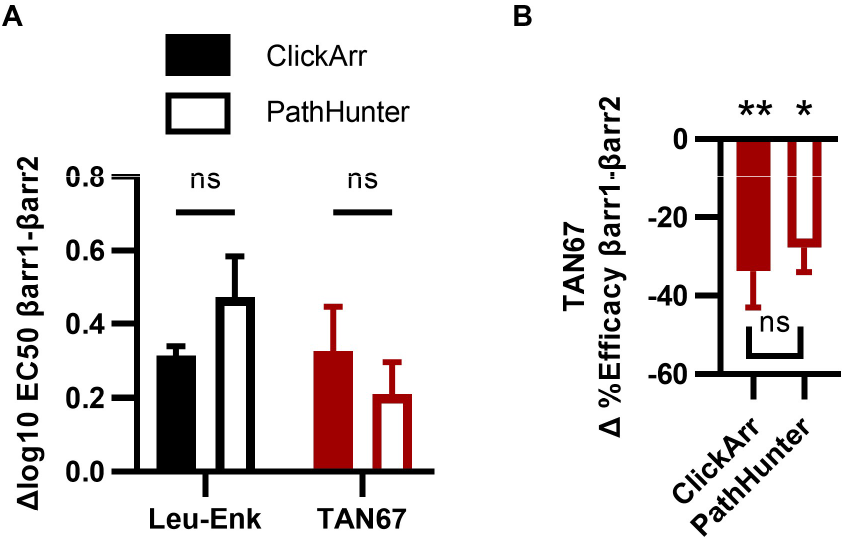
TAN67 isoform bias confirmed by PathHunter assay. A) Similar isoform potency differences are detected in both the ClickArr and the PathHunter assay. B) The PathHunter assay confirms the TAN67 efficacy bias detected in by the ClickArr assay. *, p < 0.05; **, p < 0.01 with respect to Δ%Efficacy for leu-enkephalin (defined as zero); n = 5-8.

**Supplemental Fig. 2.**
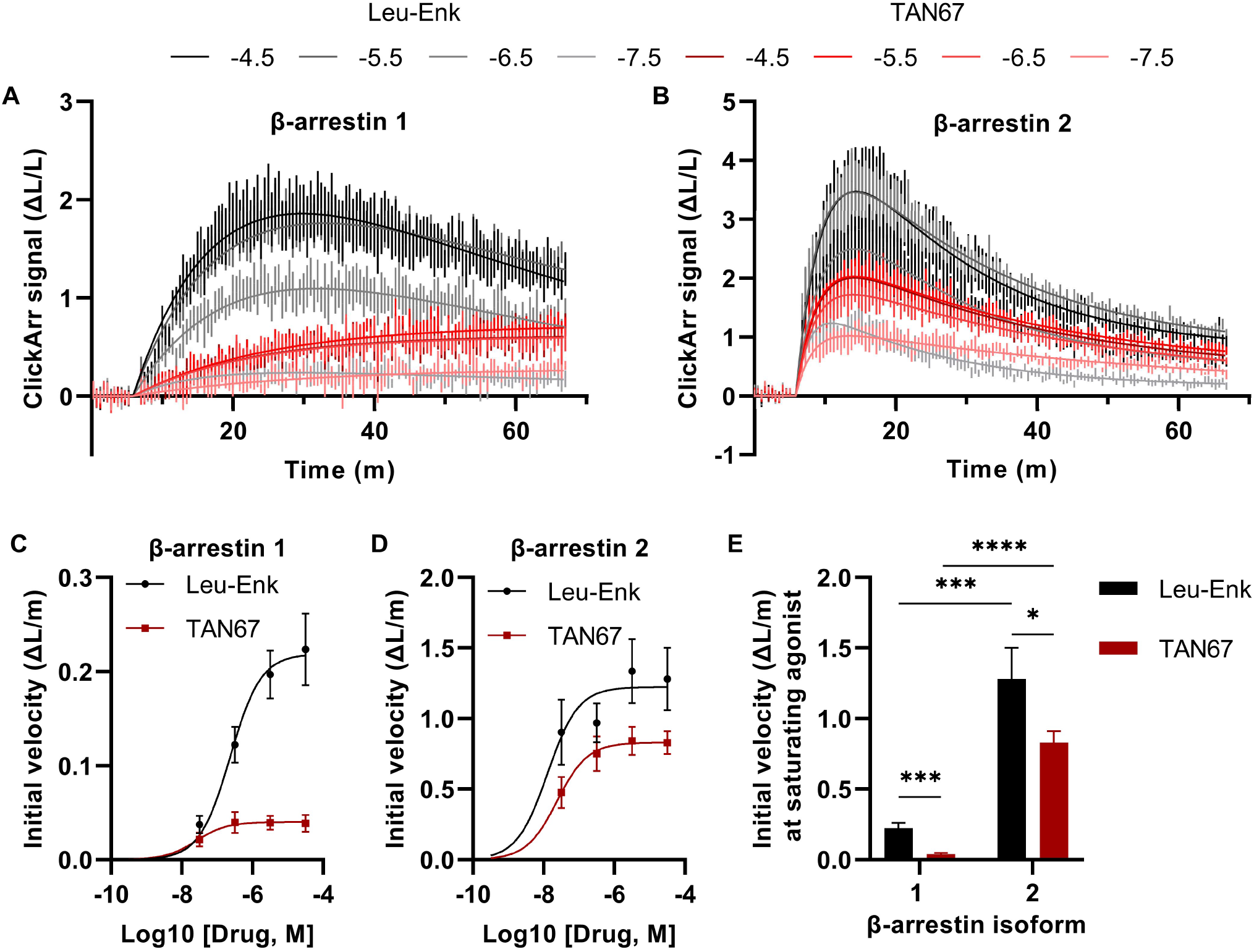
Plate format of ClickArr facilitates kinetic measurements of bias. Leu-enkephalin (Leu-Enk) and TAN67 were applied to wells containing cells co-transfected with all three ClickArr constructs to simultaneously measure kinetic profiles of four concentrations of each agonist and both β-arrestin isoforms. A,B) Arrestin recruitment signal for β-arrestin 1 (A) and β-arrestin 2 (B) over time with fitted curves (see supplemental methods). The traces in A and B were acquired concomitantly on the same plate (n = 6). The data is reported as the change in luminescence normalized to the two minutes of baseline preceding drug addition (ΔL/L), with vehicle drift subtracted. The legend is log10(concentration, M). C, D) Plotting the initial reaction velocity for β-arrestin 1 (C) and β-arrestin 2 (D) as a function of concentration confirms the relative β-arrestin isoform bias of TAN67 seen in the equilibrium experiments in Figure 1. E) Initial velocity at saturating (10^−4^·^5^ ≈ 32 μM) compound are significantly different across both drug and isoform. *, p < 0.05; **, p < 0.01; ***, p < 0.001; ****, p < 10^−4^ for indicated pair. n = 6.

